# Detecting fine and elaborate movements with piezo sensors, from heartbeat to the temporal organization of behavior

**DOI:** 10.1101/2020.04.03.024711

**Authors:** Maria Isabel Carreño-Muñoz, Maria Carmen Medrano, Thomas Leinekugel, Maelys Bompart, Fabienne Martins, Enejda Subashi, Franck Aby, Andreas Frick, Marc Landry, Manuel Grana, Xavier Leinekugel

**Affiliations:** Université de Bordeaux, Inserm U1215, Neurocentre Magendie, 33077 Bordeaux, France; IINS, CNRS UMR 5297, Bordeaux, France; Facultad de Informatica, University of the Basque Country UPV/EHU, Paseo Manuel Lardizabal 1, 20018 Donostia-San Sebastian, Spain

## Abstract

Behavioral phenotyping devices have been successfully used to build ethograms, but studying the temporal dynamics of individual movements during spontaneous, ongoing behavior, remains a challenge. We now report on a novel device, the Phenotypix, which consists in an open-field platform resting on highly sensitive piezoelectric (electro-mechanical) pressure-sensors, with which we could detect the slightest movements from freely moving rats and mice. The combination with video recordings and signal analysis based on time-frequency decomposition, clustering and machine learning algorithms allowed to quantify various behavioral components with unprecedented accuracy, such as individual heartbeats and breathing cycles during rest, shaking in response to pain or fear, and the dynamics of balance within individual footsteps during spontaneous locomotion. We believe that this device represents a significant progress and offers new opportunities for the awaited advance of behavioral phenotyping.

## Introduction

With the advent of molecular genetics and techniques allowing to manipulate neuronal physiology with unprecedented versatility and precision, the number of animal models is growing considerably, supporting a renewed interest for integrative physiology and behavioral phenotyping. However, the presumably limited introspection and language capabilities of laboratory animals promote the need for designing sophisticated behavioral readout of internal cognitive states.

The extent of the behavioral repertoire we can identify largely depends on the technologies available for the acquisition of relevant biological information. The development of video hardware and image processing algorithms sustains fast progress in behavioral phenotyping. Recent examples include 2D[1-3] and 3D[4, 5] video acquisition, which combined with machine learning algorithms allowed to identify a number of basic postures and dynamics of spontaneous behavior[1-3, 5], providing a new vision of the ethogram at the sub-second timescale[5]. Simultaneous video recordings from several angles, including through a transparent floor plate, could successfully identify the positions of the paws and other body parts, providing detailed information about the dynamic coordination of paws and body movement during locomotion[2, 3, 6-8]. However, most (if not all) internal movements underlying behavior, such as heartbeat, breathing, shivering, or the dynamics of weight and force balance during locomotion, remain out of reach from purely visual inspection. Breathing or heartbeat are too small to be captured by visual inspection. The forces resulting from muscle activity and balance are just not visual signals. Electromyogram or electrocardiogram, as all invasive approaches, are likely to seriously interfere with spontaneous behavior. Piezoelectric technology however offers sensors of exquisite sensitivity, which positioned below the floor plate can be used to collect the dynamics of movement with very high temporal precision in a totally non-invasive manner[9, 10]. Plates resting on piezo sensors or accelerometers have successfully been used to build automated ethograms, distinguishing various behaviors such as sleep, rest, grooming, etc…[6, 11-16]. But none of them provides detailed information about the precise dynamics and forces involved in single movements during spontaneous, ongoing behavior. We now report on a novel device, the Phenotypix, which consists in an open-field platform resting on highly sensitive piezoelectric (electro-mechanical) pressure-sensors. The combination of such electromechanical (EM) acquisition with video recordings and signal analysis based on time-frequency decomposition, clustering and machine learning algorithms, allowed us to detect and quantify various behavioral components with high accuracy, such as individual heartbeats and breathing cycles during rest, shaking in response to pain and fear, and the dynamics of balance within individual footsteps during spontaneous locomotion. We believe that this novel device represents a significant progress and offers new opportunities for the awaited advance of behavioral phenotyping.

## Results

### Resolution of individual movements during spontaneous behavior in the freely moving mouse

The behavioral phenotyping device (Phenotypix, Roddata, Bordeaux, France) was designed to transmit any pressure applied on the open field platform (35×45cm) to the underlying piezoelectric sensors, with minimal dumpening and resonance for a faithful transmission of any movement of the animal (see Online Methods for details). The output signal of the piezoelectric sensors was recorded in synchrony with the video (cf Figure 1A), but at much higher sampling rate (20kHz instead of 25 frames/s). The dynamics of animal movement can therefore be resolved with high temporal precision. As illustrated in Figure 1B, frequency decomposition (power spectral density) of the electromechanical (EM) signal retracing the spontaneous behavior of a wild-type (WT) mouse during a 1h open field session suggests that animal movements are mostly expressed at frequencies between 0 and 10Hz. A closer examination of specific behaviors such as walking, self grooming or sniffing revealed that frequency decomposition of the signal indeed showed a common expression of main temporal dynamics around 10Hz. Nevertheless, the signal amplitude and shape was different for each behavior, presumably resulting in signal harmonics in the 20-30Hz range (Figure 1 C-F). Another interesting observation is that the high frequency response of the device and sampling rate of the signal allowed to resolve individual movements within complex behaviors. As illustrated in Figure 1 C-F, frame-by-frame analysis of movements related to specific behaviors revealed that individual footsteps during locomotion, paw movements during self grooming of the nose, body twitches during grooming of the back, or coordinated head and nose movements during sniffing, could be identified and quantified, providing the time course and amplitude of individual movements within complex behaviors. This may be of interest in specific applications such as the experimental study of self grooming behavior. One information readily obtained from the EM signal and most likely out of reach with visual inspection, is the strength or amplitude involved in individual movements such as back or belly grooming. Recent work suggests that *Fmr1*-KO mice, a model of Fragile X syndrome, express subtle changes in grooming behavior as a sign of stress when exposed to a novel environment[17]. As illustrated in Figure S1, we have here quantified the amplitude and frequency of the EM signal associated with grooming of the back and belly in *Fmr1*-KO mice (n= 6 and 7 animals, respectively), and found that they were differentially affected. The main change regarding grooming of the back was an increase in frequency (z=-19.338, p<0.001) while amplitude was hardly affected (WT vs Fmr1-KO, z=2,193 p=0,028). On the other hand, the main change regarding grooming of the belly was an increase in amplitude (WT vs Fmr1-KO, z=29.147, p<0.001) while the frequency was hardly affected (z= −5.080, p<0.001).

**Figure 1:**
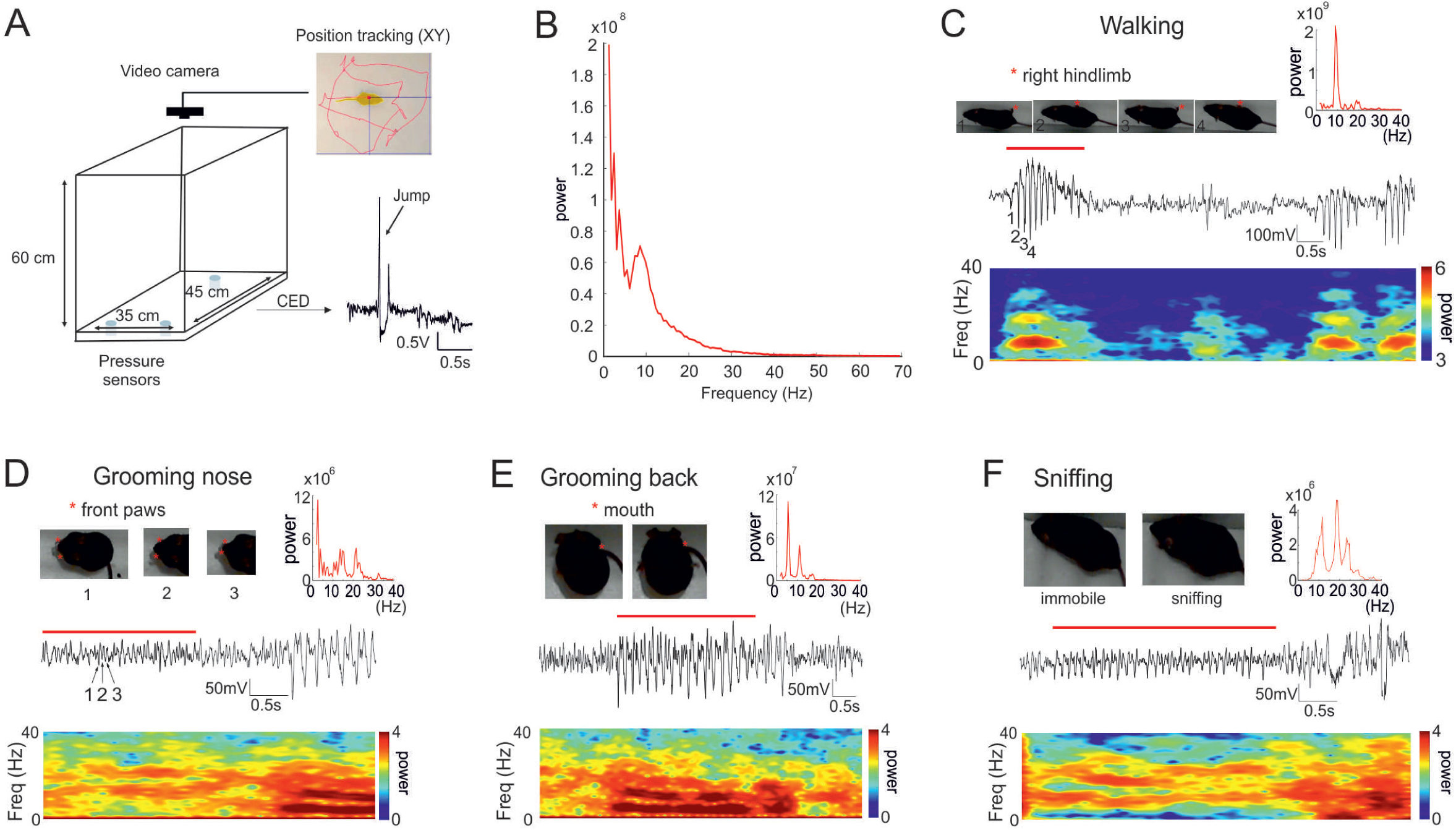
Pressure-sensor-derived detection of various body movements with the Phenotypix system. The Phenotypix system is an open field platform lying on pressure sensors allowing to detect various body movements of a freely moving rat or mouse with high sensitivity and fine time resolution. The video signal (25 frames per second) is recorded simultaneously with the digitized (20kHz) electromechanical signal from the pressure sensors, allowing synchronized offline replay for analysis. **A**. Schema of the Phenotypix acquisition system and the sensor-derived electro-mechanical (EM) signal elicited by a jumping mouse. Example tracking of the mouse with Noldus Ethovision software: yellow shape, detected mouse; red dot, central body point; vertical and horizontal blue lines, XY coordinates; red curve, mouse trajectory. **B**. Spectral composition (power spectral density) of the EM signal recorded during l h of free open field exploration. **C-F**. Typical EM signal generated by different behaviors. Each trace is shown with the corresponding time frequency spectrogram. Time of occurence of specific behaviors **(C**, locomotion> 13 cm/s; D, nose grooming, each cycle of front paw movement appears as a separate deflection in the mechanical signal; **E**, grooming of the back; **F**, sniffing, each deflection corresponds to an individual nose-movement) is indicated with an horizontal red line, illustrated with pictures taken from the video signal, and decomposed as power spectral density (PSD). Note the main component around l0Hz (and presumed harmonics in the 20-30Hz range) during active behavior.

### Breathing and heartbeat

Other very subtle movements hardly detectable from video recordings, that can also serve as an index of emotional reaction, are breathing and heartbeat. Within the EM signal obtained from a rat during sleep and immobility, we actually noticed events that seemed to correspond to breathing and heartbeat (Figure S2). The signal / noise ratio was highest during sleep and lower during rest, probably because the movements issued from the heart and chest were transferred less directly to the sensors when the animal was resting on his paws than when his chest was in direct contact with the floor-plate. As described in the literature, we did observe more regular breathing during slow wave sleep (SWS) than during REM sleep, the two main brain states here identified by the theta/delta ratio of the EEG, simultaneously recorded from the hippocampus. But more direct evidence was provided by concomitant recording of heart activity using invasive electrocardiogram (ECG) monitoring under urethane anesthesia. As illustrated in Figure 2A-B, in addition to the large and slow signal related to breathing movement, fast events occurring at about 10Hz were exactly concomitant with individual heartbeats readily visible in the ECG of both rats and mice. Taking advantage of the possibility to monitor heartbeat and breathing in a totally non invasive manner, we compared the EM signal obtained from WT freely moving mice before and after contextual fear conditioning (n = 4 and 3 animals, respectively), and as expected from the literature, we did observe statistically significant increases in breathing rate (t(5)=-17.96; p<0.001), and heart rate (t(5)=-8.42; p<0.001) after contextual fear conditioning (Figure 2C-E).

**Figure 2:**
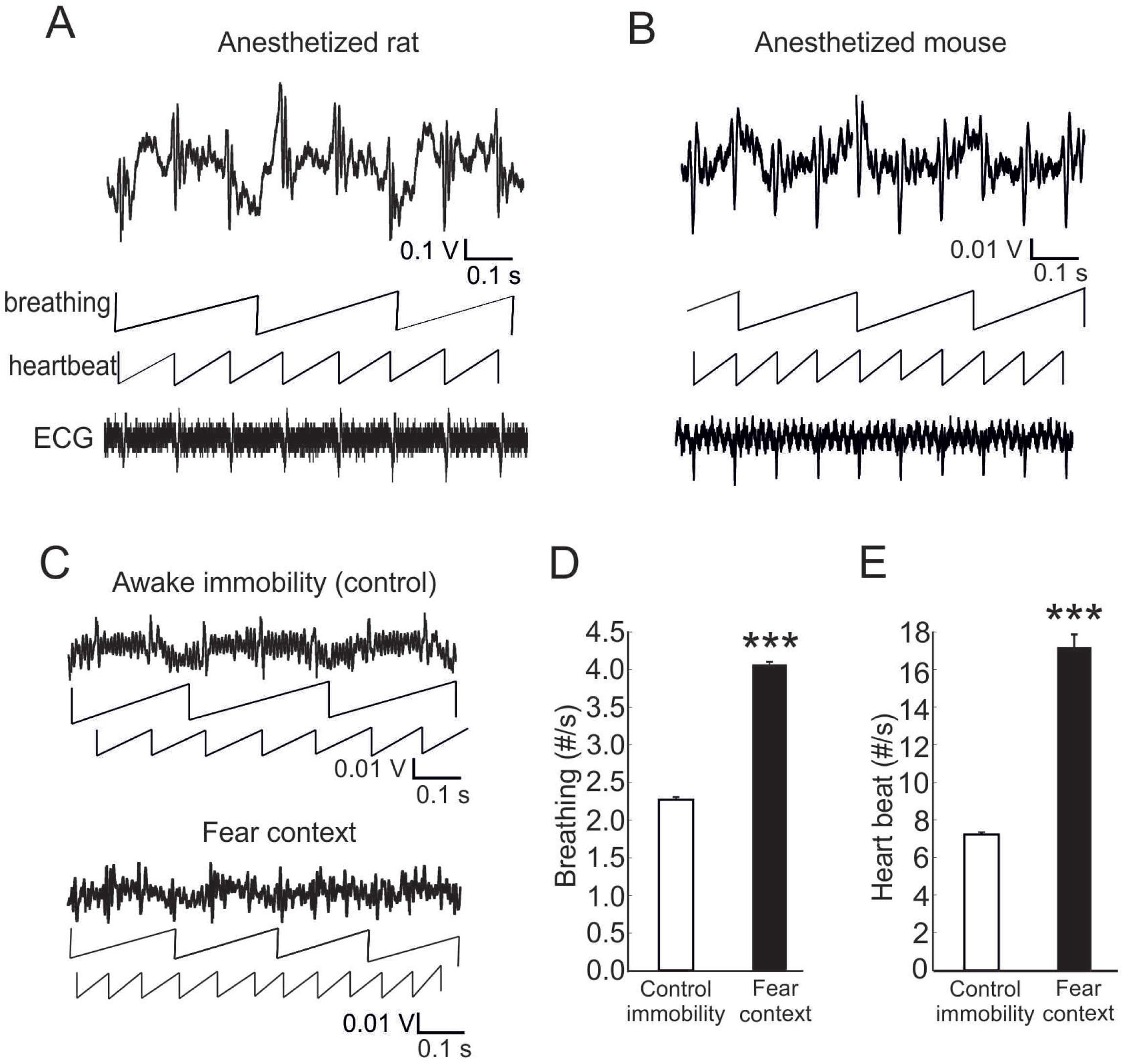
Pressure-sensor-derived detection of breathing and heart-beat. **A-B**. Simultaneously recorded electromechanical signal (upper traces) and electro-cardiogram (ECG, lower traces) from the anesthetized rat **(A)** and mouse **(B)**. Note the slow and higher frequency components of the sensor-derived signal, represented in middle traces (upper middle, linearized breathing cycles, lower middle, linearized heart-beat cycles) as broken lines. **C**. Electromechanical signal and linearized breathing and heart-beat cycles from a mouse during spontaneous immobility in a neutral context (awake immobility, control) and after fear conditioning (Fear context). **D-E**. Average breathing **(D)** and heart rate **(E)** during awake immobility in control and after fear conditioning(***, p<0.001).

### High-frequency shivering associated with pain and fear

Although the temporal dynamics of normal movements seem to be mostly confined to frequencies within the 0-10Hz range, we did notice faster components in specific experimental conditions. Immediately after surgical intervention of moderate severity (craniotomy) performed in rats without analgesic pre-medication (n=4 animals), we could feel from direct handling contact that the animal was shaking/shivering upon recovery from anesthesia, which translated in the Phenotypix as high amplitude intermittent events in the 10-45Hz frequency range (Figure 3A, left). These events were efficiently suppressed a few minutes after injection of the analgesic buprenorphine (t(5)=-11.604, p<0.001; Figure 3A, right), suggesting that they were a reaction to post-operative pain. Shaking/shivering was not observed in response to local inflammation produced by CFA injection in a rear paw (data not shown), suggesting that shaking is rather a signature of generalized pain. We did not detect either shaking in mice after surgery performed in similar conditions (n=4 animals), suggesting that rats and mice do not fully share behavioral responses to pain.

**Figure 3:**
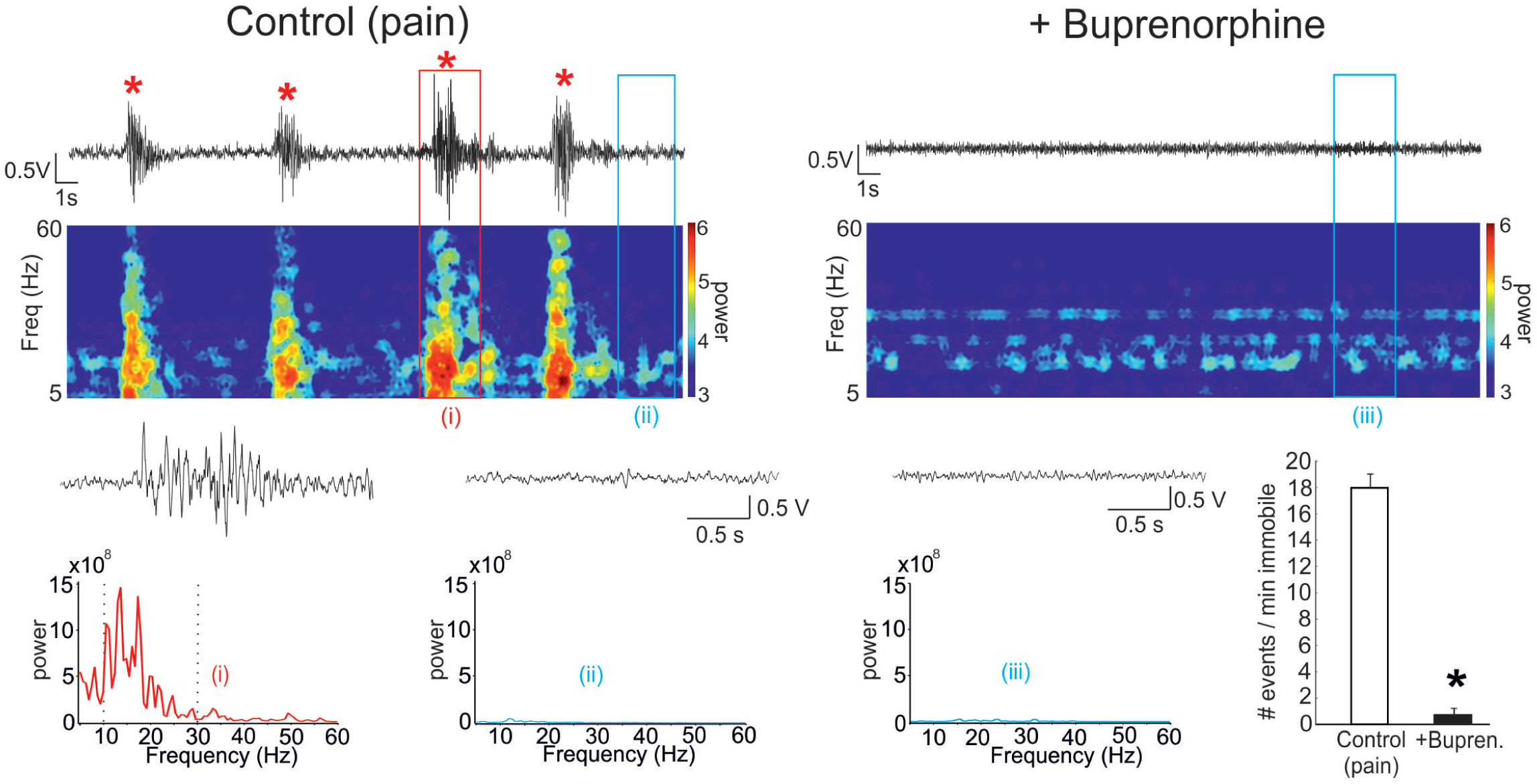
Pressure-sensor-derived detection of low frequency shaking (10-45Hz) Electromechanical signal (**top traces**, raw data) and associated time-frequency spectrogram (**underneath color plots**, left scale is frequency range, color scale indicates power) show the presence of high-power transient 10-30Hz oscillations **(left**, *) during behavioral rest in a rat upon waking up from anesthezia after surgery, that are not expressed after a pharmacological treatment against pain **(right**,**+ Buprenorphine)**. A single shaking event **(red box, i)** and a section of equivalent duration out of shaking events **(blue box, ii)**, are displayed below at wider time scale, above their corresponding Power Density Spectra. Note the high power in the 10-30Hz frequency range during pain-related shaking. The **right bar plot** shows the comparison in frequency of occurrence of 10-30Hz shaking events (detected as supra-threshold power in the 10-45Hz frequency range during immobility periods) between Control and Buprenorphine post-operative conditions.

Another behavioral condition of interest in terms of motor expression is that of fear conditioning, characterized by the active suppression of movement (freezing). While freezing behavior is classically quantified manually from the video recording, we looked for its potential specific signature in the time-frequency composition of the EM signal of mice after contextual fear conditioning. Episodes of total immobility could indeed be detected with high efficiency and reliability using a script collecting periods of EM signal below a manually set threshold of power in the 5-130Hz frequency range (Figure 4A). In addition, we noticed the high incidence of high frequency (80-120Hz) shaking events when the animal was inserted in the recording arena after fear conditioning (Figure 4B). When we compared the times of occurrence of shaking and freezing in different behavioral conditions (Figure 4C-F, n=7 animals), we found that shaking was predominantly expressed as a behavioral response to context but not to the conditioned stimulus (Control vs Fear Context, F(2,18)= 35.639, p<0.001; Control vs CS, F(2,18)= 35.639, p= 0.953), while it was the opposite for freezing (Control vs Fear context, F(2,18)=24.493, p=0.614; Control vs CS, F(2,18)= 24.493, p<0.001), raising the possibility that shaking may be a behavioral response to diffuse threat and freezing to imminent threat. A confounding factor with freezing is that it was difficult from visual inspection to distinguish between immobility periods due to the real expression of fear from behavioral immobility associated with brief rest or active scanning of the environment. Short immobility periods (<2s) were detected by our algorithm during the first minutes of exploration. In the literature, a classical approximation is to consider immobility periods longer than 2s as freezing and to ignore those of shorter durations. Accordingly, mice inserted in a novel environment (n=7 animals) expressed shaking during the first few minutes while few freezing behavior (>2s) could be identified. Nevertheless, pre-treatment with the anxiolytic Diazepam (at the non-sedative dose of 1.5mg/kg, n=6 animals) fully abolished the shaking events expressed during the first few minutes in the environment (Control vs DZ, t(11)=5.052, p<0.001), but did not significantly reduce the number of freezing episodes (Control vs DZ, t(11)=-0.517, p=0.6153). Further investigation may evaluate whether these brief immobility periods are related with mild anxiety and increased attention to potential alerts in an unfamiliar environment. The possibility that shaking and freezing might be distinct signatures of fear was further suggested by the behavioral reaction of mice exposed to the presence of a rat, one of their natural predators, which induced the immediate and remarkable expression of shaking, while inducing little freezing (Figure S3). Therefore, the Phenotypix proved efficient to detect and quantify internal movements such as heartbeat or shaking/shivering, virtually undetectable to the eye of even an experienced human observer.

**Figure 4:**
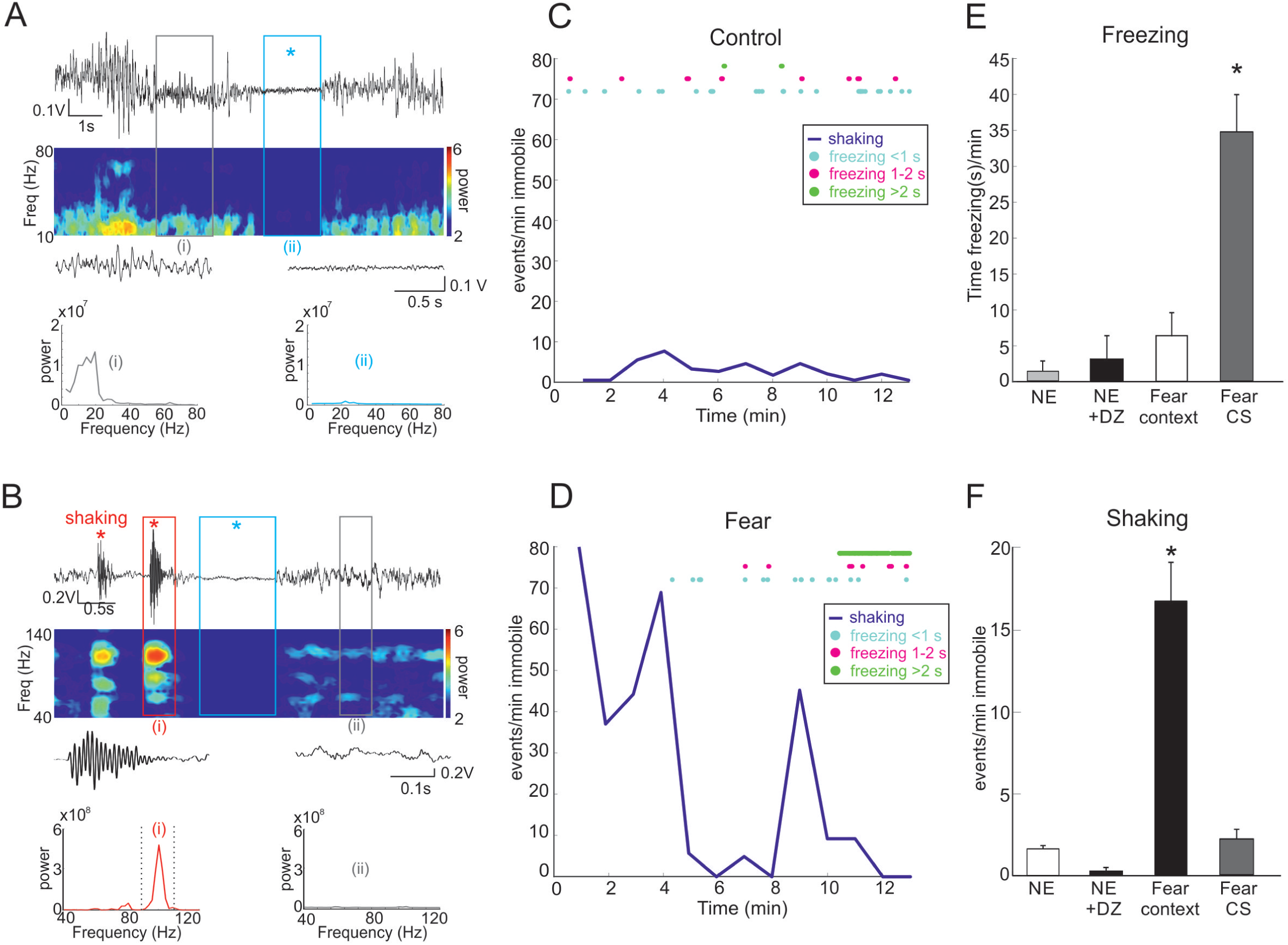
Pressure-sensor-derived detection of fear-related freezing and shaking. **A**. Electromechanical signal **(top trace**, raw data) and associated time-frequency spectrogram **(underneath color plot**, left scale is frequency range, color scale indicates power) show the presence of transient immobility periods (freezing, blue *) during spontaneous exploration of a novel environment in a naive mouse. Below are shown a single freezing event **(blue box**, ii) vs basal resting condition **(grey box**, i), displayed at wider time scale above the corresponding Power Density Spectra. Note the strong drop in power at all frequencies during freezing. **B**. Electromechanical signal **(top trace**, raw data) and associated time-frequency spectrogram **(underneath color plot**, left scale is frequency range, color scale indicates power) show the presence of spontaneous transient 80-120Hz oscillations in the signal (shaking, red*) during behavioural rest from a mouse after contextual fear conditioning. Below are shown a single shaking event **(red box**, i) vs basal resting condition **(grey box**, ii), displayed at wider time scale above the corresponding Power Density Spectra. The signal in the blue box (blue*) corresponds to a freezing episode. Note the high power in the 80-120Hz frequency range during shaking. **C-D**. Time course of occurrence of freezing (color coded **circles)**, detected as infra-threshold power in the 5-l 30Hz frequency range) and shaking **(blue line**, detected as supra-threshold power in the 65-130Hz frequency range) events over 13min of recording before **(C, Control)** and after **(D, Fear)** contextual fear conditioning. Freezing episodes are indicated according to their duration: brief (<1 s, in **cyan)**, intermediate (1-2s, in **magenta)**, long (>2s, in **green)**. **E-F. Bar plots** of global expression of spontaneous freezing **(E)** and 65-130Hz shaking events **(F)** either in control conditions (novel environment, NE), before and after pre-treatment with the anxiolytic Diazepam (+DZ), or after contextual fear conditioning **(Fear context)**. Note that while freezing is the dominant behavioural expression of fear in response to CS, shaking is the dominant expression of context-related fear. ***p<0.001.

### Gait and time course of individual footsteps

Another interesting aspect of pressure sensors is the possibility to evaluate the dynamics of the coordination and strengths of limb movements involved in locomotion. Locomotion has been extensively studied using various experimental paradigms associated with image processing tools allowing to gather increasingly sophisticated spatio-temporal information about stride or stance. Nevertheless, it has remained quite out of reach to get non invasive information about the dynamics of strengths, which can not be evaluated by visual inspection, however sophisticated. We observed that the EM signal provides some information about the dynamics of locomotion.

Frame by frame analysis of the synchronously recorded video signal allows to depict the EM signature of individual footsteps. In the short sample of spontaneous locomotion illustrated in Figure 5A, one can distinguish a few initial footsteps of small amplitude, that correspond to orienting behavior (the mouse was changing direction but not moving forward). The EM signature of locomotion, with the animal really starting to move ahead, then becomes much more visible, as series of 5 to 10 footsteps of increasing and then decreasing amplitude, displaying a spindle pattern that turned out to be very typical of mouse locomotion. Because the EM signal is the result of the dynamic distribution of weight and of all the forces generated by a multitude of muscles within the animal’s body, it is a complex mixture that depends on the coordination of the various limbs and strengths involved in movement. Nevertheless, we think that the very stereotypical signature of spontaneous locomotion in WT mice may serve as a reference template for the detection of motor impairment in various models of pathology.

**Figure 5:**
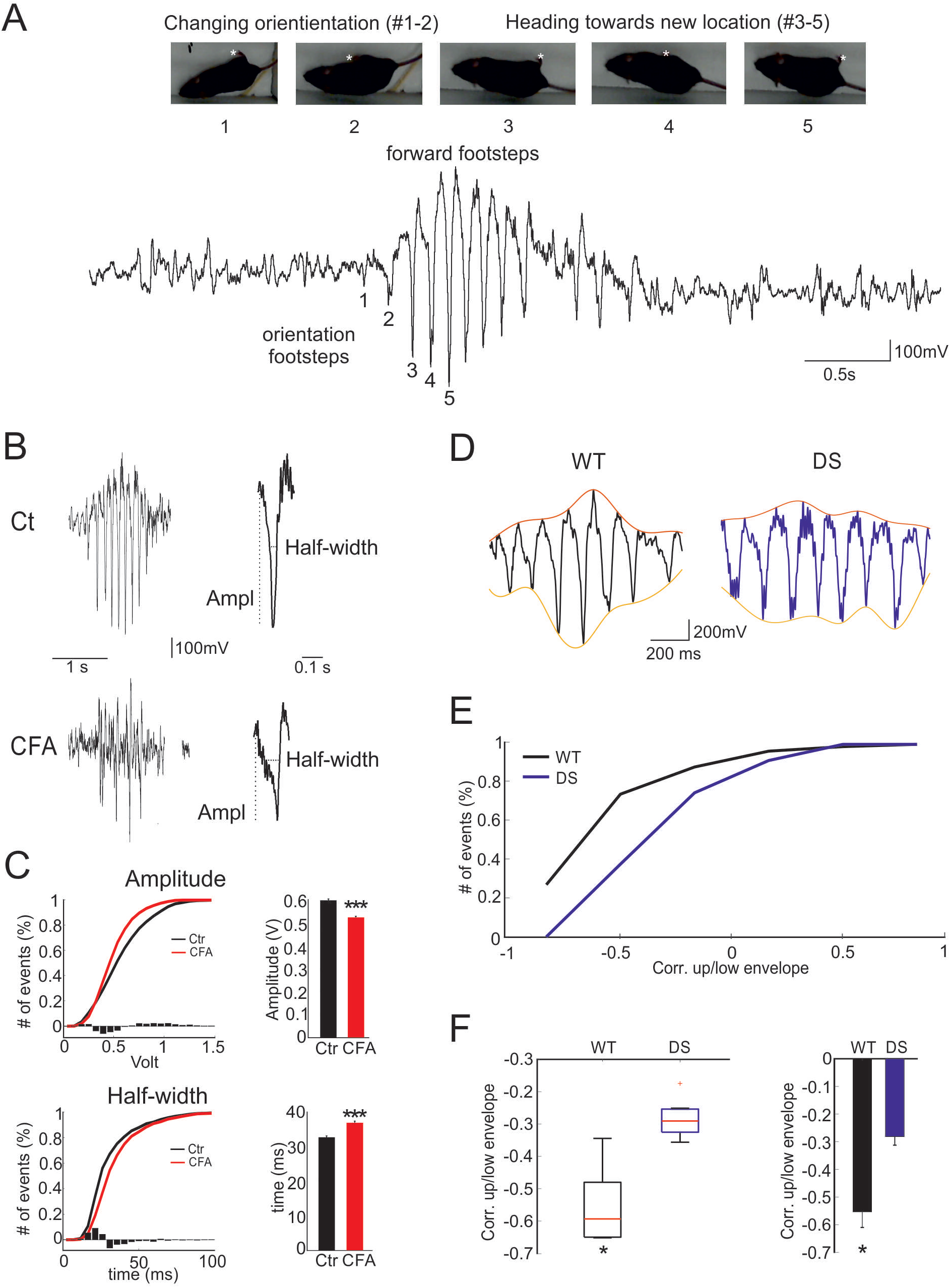
Pressure-sensor-derived signature of gait during spontaneous locomotion. **A**. Typical Electromechanical signature of mouse locomotion. Images 1-6, successive video frames(* right rear paw). Note the very light electromechanical signal generated by orientation footsteps **(#1-2**, when the mouse is changing orientation) compared to the strong signature of forward footsteps **(#3-5**, each footstep appears as a fast downward/upward deflection when the mouse is giving impulsion to move forward). **B-C**. The signature of individual footsteps during locomotion is affected in mice with a sore paw. Electromechanical signal typical ofa locomotor episode (speed 13-23 cm/s; right, zoom on an individual footstep), either in normal conditions **(Ct, upper traces)** or after CFA-induced inflammatory pain in the left rear paw **(CFA, lower traces)**. The normalized cumulative distributions **(C, left; black curves**, Ct; **red curves**, CFA; **superimposed bar histograms**, difference between Ct and CFA) and averages across animals **(C, right)** of footsteps amplitude **(upper plots)** and half-width **(lower plots)** indicate the presence of footsteps of statistically significant(***, p < 0.001) lower amplitude and slower kinetics in CFA compared to control mice. **D-F**. The correlation coefficient between the upper and lower EM signal envelopes (D, **orange and yellow lines respectively)** was computed as an index of the time course profile of successive footsteps during locomotor events (homogenous speed range, 13-30 cm/s) in control **(WT)** and Ts65Dn **(DS)** mice. The distribution of correlation coefficients of all locomotor episodes is presented as a normalized cumulative histogram **(E)**, as a moustache plot **(F, left)** and as Mean and SEM **(F, right)**. Note the statiscally different(*, p<0.05) correlation coefficients between upper and lower envelopes in WT vs DS mice.

As a first example, we recorded the EM signal of mice injected with CFA in one of the rear paws, producing local inflammation so that the animal tended to avoid pressing on the sore paw. Using a machine learning approach combining linear Support Vector Machine (SVM) classifier, autoregressive model (AR) feature extraction, and k-means clustering (see Online Methods), we identified clusters of footsteps that differentiated control from CFA mice. Visual inspection of the signal chunks identified by the discriminating clusters guided our attention towards the amplitude and time course of the EM signature of individual footsteps. We therefore performed a systematic quantification of amplitude and half width of all footsteps emitted by control and CFA mice during locomotion at comparable running speed (n=9 animals in each condition). As illustrated in Figure 5B-C, some footsteps of CFA mice indeed turned out to be smaller and of slower time course than those of control mice (control vs CFA: amplitude, z=7.781, p<0.001; half-width, z=-10.354, p<0.001), which reflects in the distributions of amplitude and half-width of the EM signal underlying individual footsteps during steady locomotion (15-30 cm/s). This is compatible with the likely consequence that CFA mice tend to avoid pressing on their sore paw.

Down Syndrom (DS) is also associated with motor impairment. Upon visual inspection of locomotion in Ts65Dn (a model of DS) and WT mice (n=5 animals in each group), we noticed a disruption of the typical spindle pattern characteristic of WT mice. As illustrated in Figure 5D-F, this was confirmed by the quantification of the correlation coefficients between the lower and upper envelopes of the EM signal associated with steady locomotion (13-30 cm/s), suggesting altered balance and movement coordination in DS (WT vs DS, t(8)=4.2212, p=0.0029).

## Discussion

A novel device (the Phenotypix), made of an open-field platform resting on highly sensitive piezoelectric pressure sensors, provided access in a totally non invasive manner to very fine components of rat or mouse spontaneous behavior. Existing systems[11-13] based on similar principles, combined with spectral decomposition and automatic classification, are used to generate ethograms, attributing each time bin of the recording to the most likely ongoing behavior, such as walking, eating, drinking, seizures, etc… But in contrast to these systems, the fine sensitivity and high sampling rate of the Phenotypix, combined with highly efficient antivibration system to minimize dumpening and resonance, allow to resolve individual movements such as individual breathing cycles, heartbeats or single footsteps during locomotion. Through the study of various behavioral conditions and transgenic models, we could identify and quantify novel behavioral components that can be useful for the study of several fields of behavioral neuroscience such as sleep, stress, pain, motor symptoms of neurodevelopmental diseases and locomotion.

Although existing devices were shown to have good performance for the detection of various kinds of self-grooming behavior[11, 13], they are not used, to our knowledge, for the quantification of the frequency and of the strength or amplitude of individual self-grooming body movements. The observation of increased amplitude and frequency of the EM signal underlying body movement in self-grooming of the back and belly in *Fmr1*-KO mice is an interesting complement to recent studies pointing at fine alterations in self-grooming behavior in the mouse under stressful conditions[18-20], because it may help better understand repetitive and self-injury behavior in FXS/ASD patients[17, 21-23].

The direct and non-invasive evaluation of breathing and heart rate may prove useful for the study of sleep apneas, a pathological condition we still poorly understand. A previous report[10] described the use of piezo sensors to detect breathing movements and heartbeats of a mouse placed on a small platform (7×13cm). Our system reached comparable sensitivity and resolution with dimensions more compatible with the study of spontaneous behavior for mice and rats. Although the dimensions of the system described here are 45×35cm, it can be extended to larger environments by the apposition of several platforms, providing a multi-compartment environment best suited to the expression of complex behavior of both rats and mice. Breathing and heartbeat are vital parameters, but also strongly related to emotions, an aspect of behavior difficult to identify in animal studies. Anxiety is classically evaluated as the avoidance of situations of innate aversion such as exposed or bright areas (e.g. center of an open field, open arms of a maze)[24]. Fear on the other hand, a more acute and stronger behavioral reaction to perceived threat, is classically quantified as freezing immobility in rat and mouse studies. From our results, we propose high-frequency (80-120Hz) shaking as a complementary and more sensitive index of fear in the mouse, expressed during exposition to a fearful situation such as a novel environment, the presence of a predator or a context previously associated with fear conditioning.

We found shaking/shivering to be expressed also as a spontaneous behavioral signature of persistent pain in the rat, although at lower frequency (10-30Hz) than in fear. Most studies about pain in rats and mice rely on the quantification of reactive pain, such as the latency between the presentation of a painful stimulus and the retraction of the affected limb (e.g. of the paw in response to von Frey filaments, of the tail in response to local heating of the skin). But these are quite indirect models of persistent and spontaneous pain, a condition of major clinical relevance[25, 26]. A remarkable recent study proposed a “mouse grimace scale” as a standardized classification of facial expression to quantify subjective pain in response to noxious stimuli[27]. This approach has been combined with machine learning algorithms and extended to the identification of other emotional states in the head-fixed mouse[28], but requires a good visual access to the face of the animal for reliable evaluation of facial expression, which may prove difficult to obtain during spontaneous behavior. With the Phenotypix, spontaneous pain is likely easier and more reliable to detect because the measure by itself should not depend on the precise moment of estimation, nor on any specific position of the animal relative to the camera. Further investigation is needed to more precisely identify the nature and intensity of pain associated with shaking/shivering behavioral response.

Our device also allowed the detection of abnormalities in the execution of locomotion, a fundamental motor function. While a number of systems are available to measure the spatio-temporal organisation of gait, analyzing the sequence of positions of the various limbs during locomotion[2, 3, 6-8, 31], the Phenotypix allowed to reveal subtle alterations in the pressure signature of individual footsteps. This compound output is the result of complex interactions, that we can not yet dissociate, between the muscular strengths and the coordination of the individual limbs involved in each footstep. Nevertheless, we could access the time course of the impulse that corresponds to individual footsteps, and identify its reduced amplitude and slower time course in limping mice. Moreover, global analysis of the dynamics of successive footsteps revealed that mouse locomotion is typically organized as series of 5 to 10 footsteps of increasing and then decreasing amplitude, a pattern that was disrupted in a mouse model of Down Syndrome. Our system therefore allowed to detect alterations of locomotion in different mouse models, suggesting access to novel criteria for gait analysis that may shed new light in the understanding of various forms of ataxia.

## Materials and Methods

### Animals

Adult (age 2-4 months) male rats and mice of different strains were recorded:

- 11 Sprague-Dawley OFA adult rats, including 2 animals recorded during resting immobility and sleep, 2 anesthetized, 4 with post-operative pain and 3 after local injection of complete Freund’s adjuvant (CFA) in a rear paw
- 45 adult wild type mice (23 3xTg-AD-WT, 10 C57B1-6J, 12 FMR1-WT), including 27 recorded in control condition, 8 after exposure to fearful conditions, 7 after pre-treatment with Diazepam, 2 anesthetized, 4 after cranial surgery, and 3 after local injection of CFA in a rear paw
- 5 transgenic mice of the Ts65Dn line, a mouse model of Down Syndrome[32]
- 7 transgenic *Fmr1*-KO mice, deficient for both FMR1 RNA and FMR Protein, a model of Fragile X Syndrome [33]

All animals were bred in the laboratory animal facility in collective cages, and transferred to individual cages for the duration of the experiments. Animals were kept on a 12h/12h light/dark cycle, provided with nesting material and food and water ad libitum. All experiments were performed during the light period under constant mild luminosity (60-70 Lux). All experimental procedures were performed in accordance with the EU directives regarding the protection of animals used for experimental and scientific purposes (86/609/EEC and 2010/63/EU), with the French law, and approved by the Ethical committee CEEA50 (saisines #15349, 15350, 10897 and 50120156-A).

### Behavioral data acquisition

The mice were introduced individually onto the recording platform (Phenotypix, Roddata, Quinsac, France), a dimly illuminated open field environment (45×35cm), surrounded by 60cm-high walls and equipped with video monitoring. The epoxy floor-plate and the walls were sprayed and wiped clean with 70% ethanol before the introduction of each animal. Spontaneous behavior was recorded continuously for durations ranging from 5min to 3h. In this system, the floor plate is resting on 3 evenly distributed (as a triangle) piezoelectric pressure sensors, all connected together to a single charge amplifier, providing a continuous voltage analog signal (bandpass 0.1Hz - 9KHz) proportional to the pressure exerted on the sensors underlying the floor plate, so that any subtle changes in floor-plate pressure due to animal movement could readily be detected. Unlike other existing phenotyping systems based on analyzing the vibrational pattern of the floor-plate to identify ongoing behavior[11-13], the Phenotypix rather collects the minute pressor changes resulting from individual animal movements, which requires optimal signal preservation and was achieved by minimizing dumpening and resonance. For this purpose, the platform laid on an antivibration table (TMC), made of a plain stainless steel table top (weight about 150kg) resting on Gimbal Pistons using air pressure to keep the table top above a heavy (about 100kg) 4-legs frame. Lighter isolation platforms with pneumatic isolators (Newport Benchtop) did not prove efficient enough to preserve the pressor-derived signal from vibrations, and the performance of the Phenotypix were seriously degraded. Video signal was acquired at a sample rate of 25 frames/s with a webcam placed 1m above the platform, and the electromechanical signal was recorded continuously at a sampling rate of 20 kHz. Both signals were acquired synchronously using a Power1401 digitizer and Spike2 software (CED) and stored on a PC for offline analysis with EthoVision XT software (Noldus) and custom-made matlab scripts (Mathworks).

### EEG/EMG recording in freely moving rats and mice

Rats and mice were deeply anesthetized with isoflurane (2-5%) and implanted with individual 50µm-diameter insulated tungsten-wires connected to an Omnetics connector fixed to the cranial bone with dental cement. For EEG recordings, the electrodes were placed within the hippocampus under EEG monitoring, at the following coordinates (Paxinos atlas): AP −3.3, L 2.5, V −2.5 to −2.8 for rats, and AP - 1.95, L 1.35, V −1.2 to −1.4 for mice. For EMG recordings, a single wire, from which insulation was removed at the tip for about 2mm, was inserted into the neck muscle. The skin was put back into place and maintained with surgical glue (Gluture, WPI). Before recording, the animals were kept under daily monitoring for 1 week for healing and recovery. For recording, the animal was plugged to the recording system (Neuralynx L8 amplifiers) with a tethered headstage (HS-16, Neuralynx), and the wide band (0.1Hz-9kHz) digitized signal continuously recorded with CED-Spike2 (in synchrony with the behavioral data) and stored on PC for offline analysis.

### ECG recording under anesthesia

Rats and mice were deeply anesthetized with urethane (15mg/kg), placed on the recording platform, and the electrocardiogram recorded as the voltage difference between 2 electrical wires positioned subcutaneously in the upper chest and hindpaw. The wide band (0.1Hz-9kHz) digitized ECG signal was continuously recorded with CED-Spike2 (in synchrony with the behavioral data) and stored on PC for offline analysis.

### Pain

A group of rats and WT mice received a subcutaneous injection of complete Freund’s adjuvant (CFA) in a hindpaw as a model of persistant inflammatory pain[35]. Male *Sprague-Dawley* (SD) rats (180–220g) and male C57BL/6J mice (20-25g) were immobilised with a specific contortion technique to allow a good access to one of their hindpaw. A single dose (100µl in rats, 20µl in mice) of complete Freund’s adjuvant (CFA, SIGMA Aldrich) was injected subcutaneously into the plantar surface of one hind paw using a microliter syringe with a 27-gauge needle. The CFA injection immediately induces local inflammation, paw swelling and pain[35]. The animals were recorded in the Phenotypix 24 hours after the injection, for a duration of 1h. In another series of pain experiments, rats and mice acutely implanted with intracranial electrodes for electrophysiological recordings were recorded in the Phenotypix 10min after recovery from the anesthesia, for a 5 min recording session. Immediately after, these animals were injected with a single dose of the analgesic buprenorphine (0,05mg/Kg; 0.1ml/10g animal, IP), and 10 min after the injection, recorded again for 5 min.

### Fear conditioning

On day 1, WT mice were transferred from their housing room to the recording room for a fear conditioning session. After a 5 min acclimation period to the conditioning chamber, 5 trials (intertrial interval 5-10min) were performed, each consisting of 10 intermittent white tones (500ms duration separated by 1s), the last 5 of which paired with electrical footschocks. The mice were brought back to their housing room after the conditioning session. On day 2, the mice were tested for contextual fear by being inserted in the recording chamber (different from the conditioning chamber but in the same experimental room with the same contextual configuration) for 15 min of free exploration after which 3 series of tones (conditioned stimulus, CS) were presented.

### Exposure to predator

Individual WT mice could freely explore the recording chamber of the Phenotypix for 1 hour. On the next day, they were placed again in the recording chamber, from which one of the arena walls had been removed and replaced by a cage containing an adult Spague Dawley rat, with a grid separating the rat (separated compartment, not in contact with the piezo sensors) from the mouse (placed on the Phenotypix platform). The spontaneous behavior of the mouse was recorded during 15min.

### Data analysis

The piezosensor-derived signal was first explored visually together with the video for identification of spontaneous behaviours. After visualization of the raw data, manual tagging of the different behaviors was performed by a trained expert, using Spike2 software. For further processing, the raw data was downsampled from 20KHz to 1250Hz using ndmanager pluggins[36]. Sonic Visualizer software was used to explore visually the time-frequency domain of the piezo derived-signal. Running periods were selected based on the animal velocity, calculated from the XY coordinates obtained through offline automatic animal tracking with Ethovision XT software (Noldus). Grooming amplitude was quantified on manually selected periods as the peak-to-through amplitude of each body movement-related signal deflection. Shaking detection was done only during periods of immobility (speed <2.5cm/s during a least 1s). Automatic detection of shaking events was performed as threshold crossing on the bandpass filtered (10-45Hz for pain, 65-130Hz for fear), squared and normalized signal. Automatic detection of freezing events was performed as threshold crossing on the 5-130Hz bandpass filtered, squared and normalized signal. The thresholds for event detection were adjusted manually for each recording, and the outcome of the detection verified manually using Neuroscope visualization software[36]. Signal decomposition (Fourier Analysis) and quantification was performed using custom mat lab (The Math works) code, available upon request.

The dynamics of locomotion were investigated within a similar speed range in WT and DS mice, during periods of intermediate running speed, between 13 and 30 cm/s for a minimum duration of 700ms. The corresponding EM signal, downsampled to 1250Hz, was extracted using a sliding window of 800 samples. In order to discriminate between the features of locomotion signal issued from control vs CFA treated animals, we have used Support Vector Machine (SVM). The regression coefficients obtained from an autorregressive model [37] of order p were used as input features for k-means clustering [38]. We next identified “pure” clusters, containing samples from only either CFA or control mice, and performed on this sub-dataset a classification process using a linear support vector machine (SVM) [39] approach. In order to find the largest feature dataset with the highest classification performance, we conducted a grid search on the following parameters: AR order (p), window size (w) and number of clusters (k) maximizing an objective function defined by the product of the feature dataset size and the classification performance measured by F-score [40]. From the feature dataset maximizing the objective function we retrieved the corresponding signal windows, merging them to obtain the signal chunks where discrimination had been detected. Finally, we carried out the extraction of manually defined descriptive features on these signal chunks, which showed significant differences. Moreover, it is possible to build predictive models of the mice class following this approach as shown by crossvalidation experiments in which there is a separation of train and test data sets before the selection process and the construction of the SVM model, ensuring that there is no double dipping effect [41] in the estimation of the classification performance. F-score results reached 0.80 (data not shown), which is well above random choice classification.

Because visual inspection of the signal chunks identified by the discriminating clusters guided our attention towards the amplitude and time course of the EM signature of individual footsteps, we have performed a systematic quantification of amplitude and half width of all footsteps emitted by control and CFA mice during locomotion at comparable running speed (Figure 5B-C). Periods of locomotion were selected during which the animal was moving between 13 and 30cm/s without interruption and reaching at least 20cm/s. Individual footsteps were identified as consecutive suprathreshold peak-trough-peak sequences from the EM signal, bandpass filtered at various frequencies using zero-phase distorsion filters (i.e. filtering in the forward and backward direction to prevent phase-distorsion). Peaks and troughs were detected as local extremas in the 0-300Hz passband filtered EM-signal, within 50ms of either the minima detected from the 0-50Hz passband filtered signal (approximative troughs) or of the maxima detected from the 0-20Hz passband filtered EM-signal (approximative peaks), respectively. Bandpass filtered 0-5Hz signal was taken as baseline, and only local minima (troughs) of amplitude larger than 1SD from baseline were selected for further footstep analysis. The amplitude of footsteps was measured as the difference between the trough and the mean of its pre- and post-peaks. The half-width was measured as the width at half amplitude.

Locomotion and gait were also analyzed at the more global level of footsteps dynamics (Figure 5D-F) by comparing the envelopes of locomotion-related EM signal across conditions. Periods of locomotion were selected during which the animal was moving for at least 500ms between 13 and 30cm/s. The linear correlation coefficient between upper and lower envelopes was computed for each locomotor period.

### Statistics

Data processing was performed with custom-made scripts and functions from the Matlab statistics toolbox (Mathworks). Data were systematically tested for Normal distribution, either with the Lilliefors test, a modification of the Kolmogorov-Smirnov test recommended for small sample sizes [42], or with the Kolmogorov-Smirnov test for sample sizes >50. Homoscedasticity was assessed using the Levene’s test. Data following a Normal distributions and with homogenous variances were analyzed using parametric tests: ANOVA (F) with genotype or condition as factors (and post hoc Turkey test) for independent data sets, and Student t-test (t) for paired data. Data following non-normal distributions were analyzed using the non-parametric tests Kruskal-Wallis (X2) for multiple groups comparisons (posthoc, Dunn-Sidak test) or Wilcoxon ranksum test (z) for pairwise comparison of unpaired data. Results were considered significant for values of p<0.05. Data are presented as mean ± SEM.

## Data availability

The data shown in the paper will be made available upon reasonable request to the corresponding author.

## Code availability

The custom code used for analysis will be made available upon reasonable request to the corresponding author.

## Acknowledgements

We would like to thank Yannick Jeantet for fruitful discussions and comments, Delphine Gonzales, Nathalie Aubailly, and all the personnel of the Animal Facility of the NeuroCentre Magendie for animal care.

## Funding

This work was supported by funding from: INSERM (XL), Région Nouvelle Aquitaine (XL), Agence Nationale pour la Recherche (ANR, XL). We thank the Animal Housing and Genotyping facilities, supported by funding from INSERM and LabEX BRAIN (ANR-10-LABX-43). MICM was supported by an international PhD fellowship (ANR-10-IDEX-03-02). The funders had no role in study design, data collection and analysis, decision to publish, or preparation of the manuscript.

## Author contributions

XL conceived the project and collected pilot data. MICM, MCM and XL planned the research. MICM, MCM, MB, FM, ML, FA and XL participated in animal preparation. MICM, MCM, MB, FM, ES and XL performed data acquisition. AF provided lab and breeding space. MICM, MCM, MB, FM, MG, TL and XL analyzed the data. MICM, MCM, MG and XL wrote the paper.

## Potentially competing interests

XL is shareholder of Roddata. MICM, MCM, TL, MB, FM, ES, FA, AF, ML and MG declare no competing interests.

---

**Figure S1:**
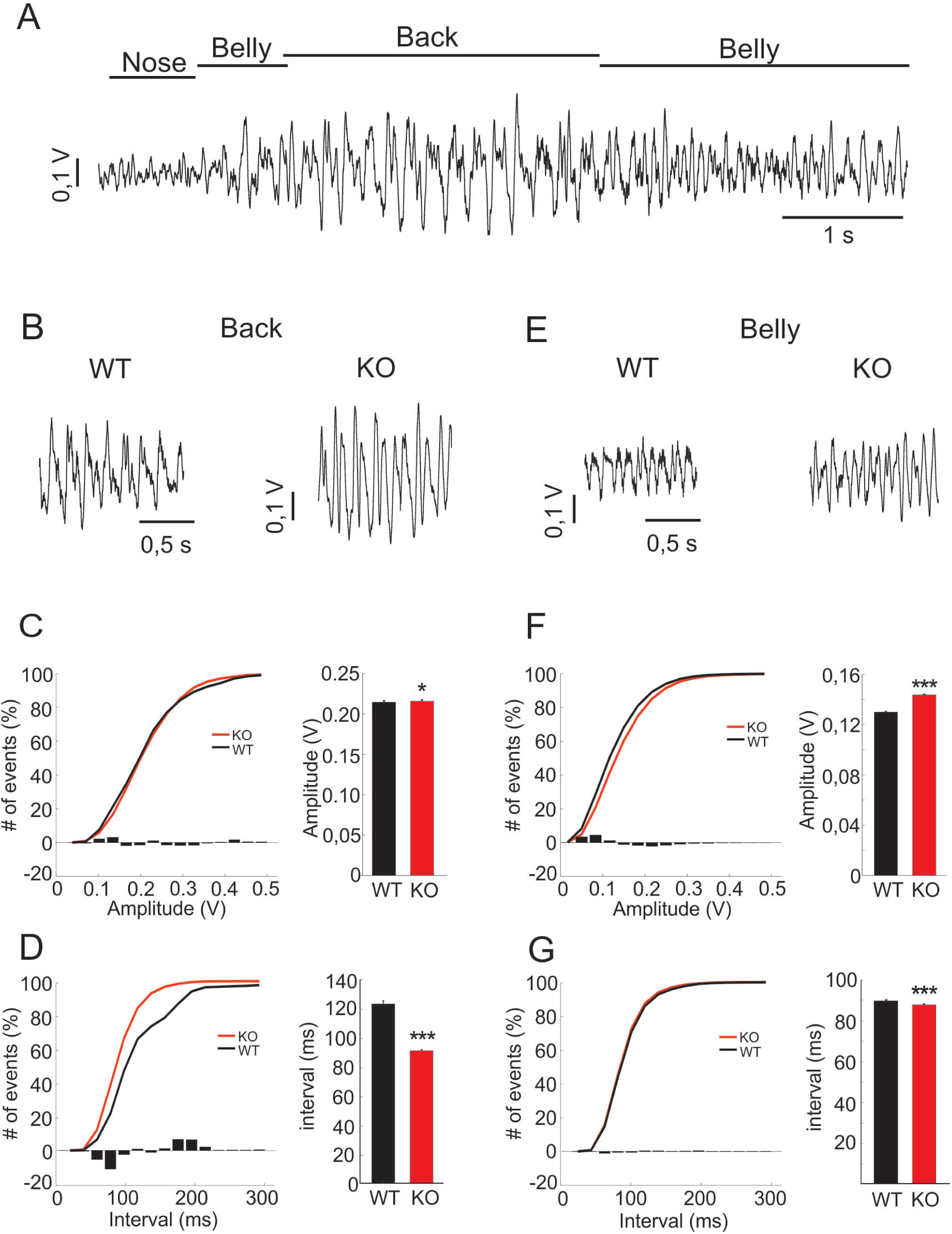
Pressure-sensor-derived signature for grooming of the back and belly in WT and *Fmr1-*KO mice. **A**. Typical sensor-derived signature of grooming the nose, back and belly in a WT mouse. **B-D**. Electromechanical signal typical of a back-grooming episode, in WT mice **(B, WT)** and *in Fmr1-*KO mice **(B, KO)**. The normalized cumulative distributions **(C** and **D, left; black curves**, WT; **red curves**, KO; superimposed bar histograms, difference between WT and KO) and averages across animals **(C** and **D, right)** of grooming signal amplitude **(C)** and inter-movement interval **(D)**. **E-G**. Same as in C-D but for grooming of the belly, indicating the amplitude and frequency of grooming of the belly *in Fmr 1-*KO compared to WT mice. Note the clear increases in amplitude for grooming of the belly and in frequency (ie decreased inter-event intervals) for grooming of the back *in Fmr1-*KO compared to WT mice(*, p < 0.05, ***, p<0.001).

**Figure S2:**
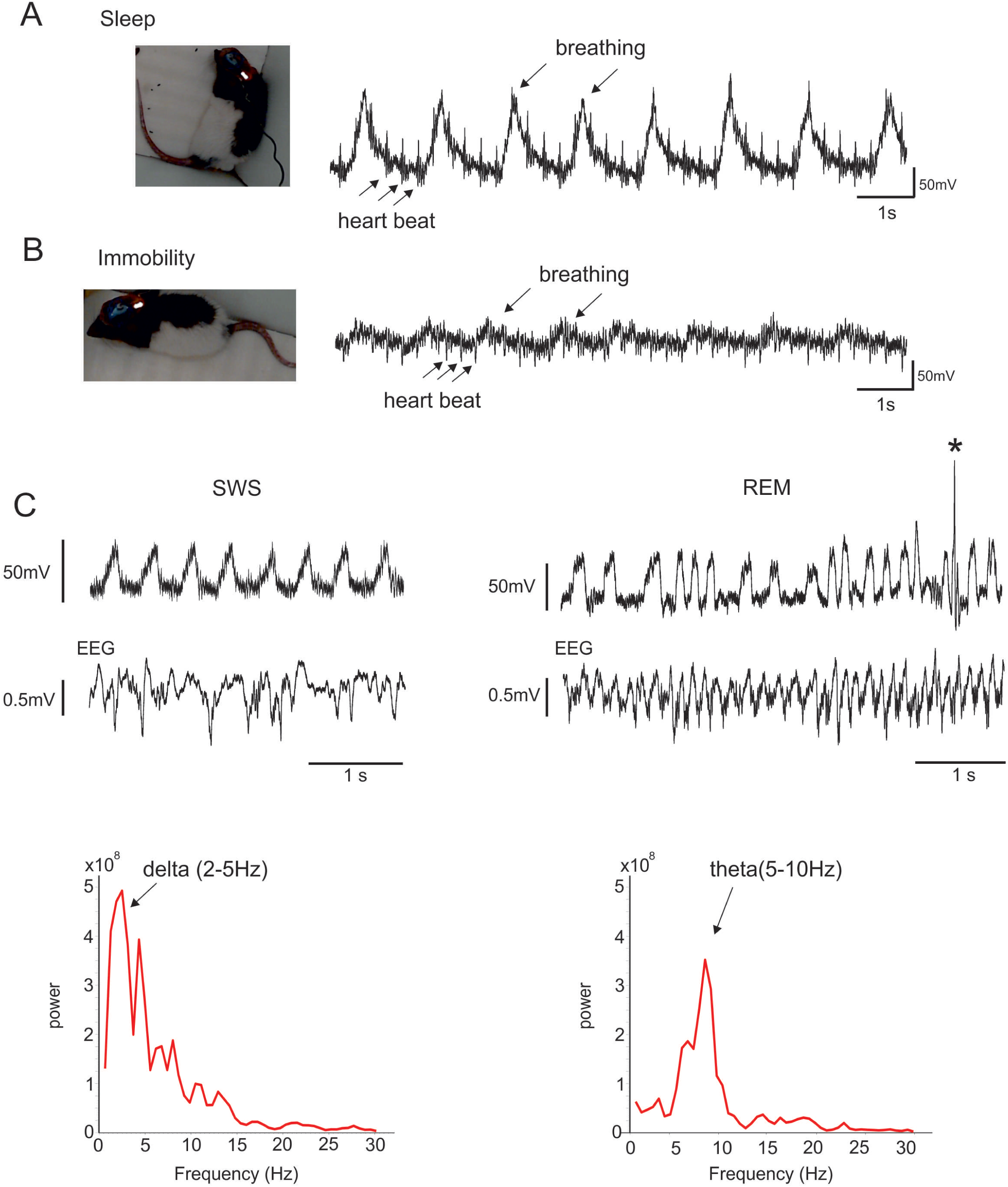
Pressure-sensor-derived detection of breathing and heart-beat during immobility and sleep. **A-B**. Sensor-derived signal from a rat during sleep and immobility. Note the higher amplitude ofbreathing and heart-beat signals when the animal is lying on the floor **(A**, Sleep) compared to resting on his paws **(B**, immobility). **C**. Simultaneously recorded sensor-derived (upper trace) and EEG (lower trace) signal from a mouse during sleep. Power spectral decomposition of EEG traces (below) identifies slow wave sleep (SWS, left traces) as high delta (2-5Hz) and low theta (6-l0Hz) power, and REM sleep (right traces) as low delta and high theta power. Note that SWS is characterized by very regular breathing and absence of any other movement, while REM sleep shows irregular breathing as well as occasional movements (twitches, *).

**Figure S3:**
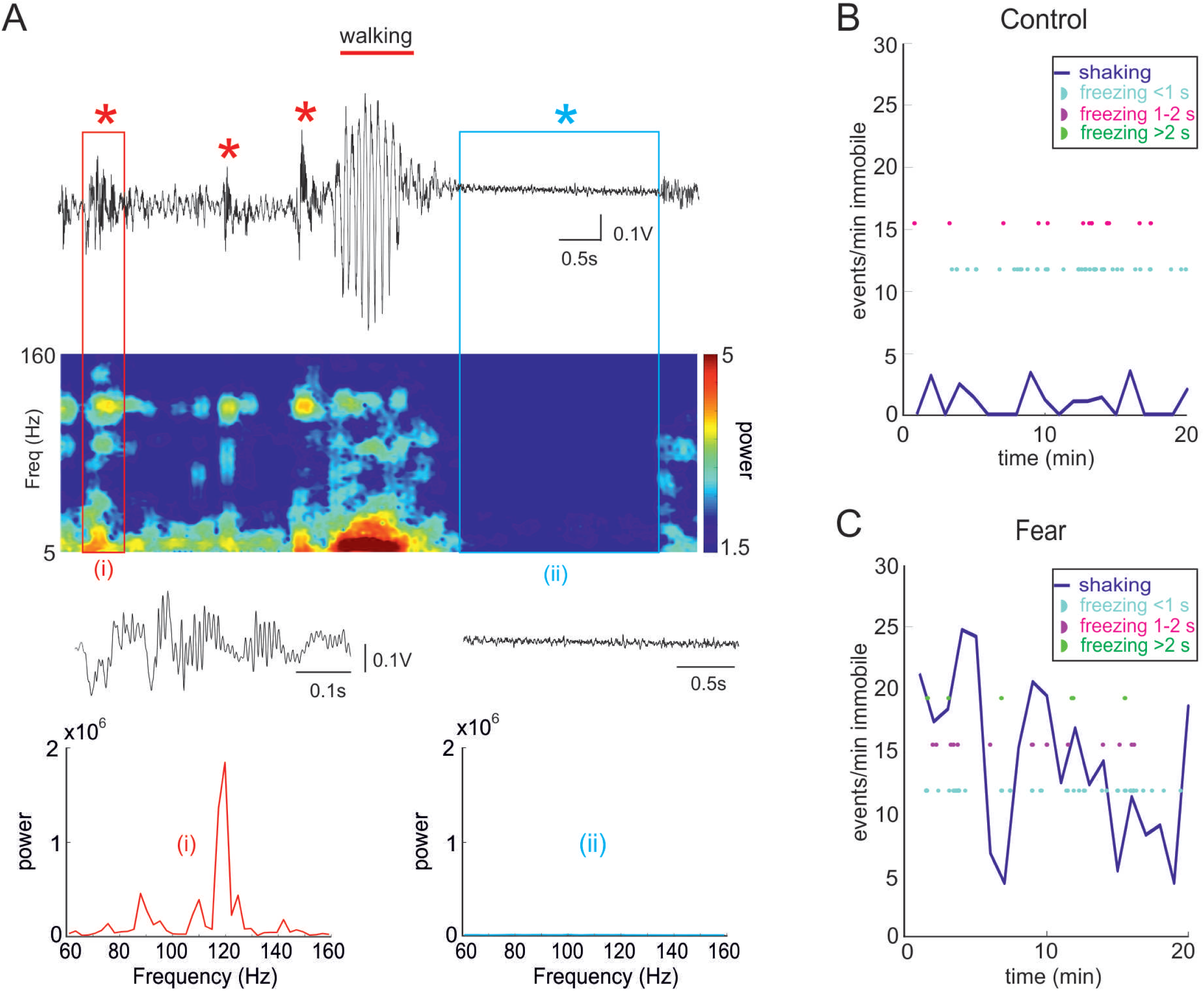
Pressure-sensor-derived detection of fear of predator. **A**. Electromechanical signal **(top trace**, raw data) and associated time-frequency spectrogram **(underneath color plot**, left scale is frequency range, color scale indicates power) show the presence of both freezing **(blue*)** and 80-130Hz shaking **(red*)** during exposure of a mouse to a predator (a rat behind bars on the side of the open field, so that the mouse can see and smell but not get in direct physical contact with it). Below are shown a single freezing event **(blue box, ii)** vs 65-130Hz shaking event **(red box, i)**, displayed at wider time scale above the corresponding Power Density Spectra. Note the high power in the 65-130Hz frequency range during shaking vs the strong drop in power at all frequencies during freezing. **B-C**. Time course of occurrence of freezing (color coded **half-circles)** and shaking **(blue line)** events over 20min before **(B, Control)** and during **(C, Fear)** exposition to the predator. Freezing episodes are indicated according to their duration: short (1-2s, in **magenta ticks)** or long (>2s, in **green ticks)**. Note that shaking is the dominant expression of fear in response to the exposure to a predator.

## References

1. Kabra, M., Robie, A.A., Rivera-Alba, M., Branson, S., and Branson, K. (2013). JAABA: interactive machine learning for automatic annotation of animal behavior. Nat.Methods 10, 64–67.

2. Pereira, T.D., Aldarondo, D.E., Willmore, L., Kislin, M., Wang, S.S.H., Murthy, M., and Shaevitz, J.W. (2019). Fast animal pose estimation using deep neural networks. Nature Methods 16, 117–125.

3. Mathis, A., Mamidanna, P., Cury, K.M., Abe, T., Murthy, V.N., Mathis, M.W., and Bethge, M. (2018). DeepLabCut: markerless pose estimation of user-defined body parts with deep learning. Nature Neuroscience 21, 1281–1289.

4. Ou-Yang, T.H., Tsai, M.L., Yen, C.T., and Lin, T.T. (2011). An infrared range camera-based approach for three-dimensional locomotion tracking and pose reconstruction in a rodent. J Neurosci.Methods 201, 116–123.

5. Wiltschko, A.B., Johnson, M.J., Iurilli, G., Peterson, R.E., Katon, J.M., Pashkovski, S.L., Abraira, V.E., Adams, R.P., and Datta, S.R. (2015). Mapping Sub-Second Structure in Mouse Behavior. Neuron 88, 1121–1135.

6. Clarke, K.A., and Still, J. (1999). Gait analysis in the mouse. Physiology & behavior 66, 723–729.

7. Machado, A.S., Darmohray, D.M., Fayad, J., Marques, H.G., and Carey, M.R. (2015). A quantitative framework for whole-body coordination reveals specific deficits in freely walking ataxic mice. eLife 4, e07892.

8. Zorner, B., Filli, L., Starkey, M.L., Gonzenbach, R., Kasper, H., Rothlisberger, M., Bolliger, M., and Schwab, M.E. (2010). Profiling locomotor recovery: comprehensive quantification of impairments after CNS damage in rodents. Nat Meth 7, 701–708.

9. Sato, S., Yamada, K., and Inagaki, N. (2006). System for simultaneously monitoring heart and breathing rate in mice using a piezoelectric transducer. Med Biol Eng Comput 44, 353–362.

10. Sato, and Shinichi (2008). Quantitative evaluation of ontogenetic change in heart rate and its autonomic regulation in newborn mice with the use of a noninvasive piezoelectric sensor. American Journal of Physiology - Heart and Circulatory Physiology 294, H1708–H1715.

11. Brodkin, J., Frank, D., Grippo, R., Hausfater, M., Gulinello, M., Achterholt, N., and Gutzen, C. (2014). Validation and implementation of a novel high-throughput behavioral phenotyping instrument for mice. J.Neurosci.Methods 224, 48–57.

12. Wood, N.I., Goodman, A.O.G., van der Burg, J.M.M., Gazeau, V., Brundin, P., Björkqvist, M., Petersén, Å., Tabrizi, S.J., Barker, R.A., and Jennifer Morton, A. (2008). Increased thirst and drinking in Huntington’s disease and the R6/2 mouse. Brain Research Bulletin 76, 70–79.

13. Chen, S.-K., Tvrdik, P., Peden, E., Cho, S., Wu, S., Spangrude, G., and Capecchi, M.R. (2010). Hematopoietic Origin of Pathological Grooming in Hoxb8 Mutant Mice. Cell 141, 775–785.

14. Mang, G.M., Nicod, J., Emmenegger, Y., Donohue, K.D., O’Hara, B.F., and Franken, P. (2014). Evaluation of a piezoelectric system as an alternative to electroencephalogram/electromyogram recordings in mouse sleep studies. Sleep 37, 1383–1392.

15. Donohue, K.D., Medonza, D.C., Crane, E.R., and O’Hara, B.F. (2008). Assessment of a non-invasive high-throughput classifier for behaviours associated with sleep and wake in mice. Biomed.Eng Online. 7, 14.

16. Daldrup, T., Remmes, J., Lesting, J., Gaburro, S., Fendt, M., Meuth, P., Kloke, V., Pape, H.C., and Seidenbecher, T. (2015). Expression of freezing and fear-potentiated startle during sustained fear in mice. Genes, Brain and Behavior 14, 281–291.

17. Carreno-Munoz, M.I., Martins, F., Medrano, M.C., Aloisi, E., Pietropaolo, S., Dechaud, C., Subashi, E., Bony, G., Ginger, M., Moujahid, A., et al. (2018). Potential Involvement of Impaired BKCa Channel Function in Sensory Defensiveness and Some Behavioral Disturbances Induced by Unfamiliar Environment in a Mouse Model of Fragile X Syndrome. Neuropsychopharmacology 43, 492–502.

18. Song, C., Berridge, K.C., and Kalueff, A.V. (2016). ’Stressing’ rodent self-grooming for neuroscience research. Nat Rev Neurosci 17, 591–591.

19. van Erp, A.M.M., Kruk, M.R., Meelis, W., and Willekens-Bramer, D.C. (1994). Effect of environmental stressors on time course, variability and form of self-grooming in the rat: Handling, social contact, defeat, novelty, restraint and fur moistening. Behavioural Brain Research 65, 47–55.

20. Meshalkina, D.A., and Kalueff, A.V. (2016). Commentary: Ethological Evaluation of the Effects of Social Defeat Stress in Mice: Beyond the Social Interaction Ratio. Frontiers in Behavioral Neuroscience 10.

21. Baranek, G.T., Foster, L.G., and Berkson, G. (1997). Tactile Defensiveness and Stereotyped Behaviors. American Journal of Occupational Therapy 51, 91–95.

22. Merenstein, S.A., Sobesky, W.E., Taylor, A.K., Riddle, J.E., Tran, H.X., and Hagerman, R.J. (1996). Molecular-clinical correlations in males with an expanded FMR1 mutation. American Journal of Medical Genetics 64, 388–394.

23. Symons, F.J., Clark, R.D., Hatton, D.D., Skinner, M., and Bailey, D.B. (2003). Self-injurious behavior in young boys with fragile X syndrome. American Journal of Medical Genetics Part A 118A, 115–121.

24. Prut, L., and Belzung, C. (2003). The open field as a paradigm to measure the effects of drugs on anxiety-like behaviors: a review. Eur J Pharmacol 463, 3–33.

25. Mogil, J.S., Davis, K.D., and Derbyshire, S.W. (2010). The necessity of animal models in pain research. Pain 151, 12–17.

26. Mogil, J.S., and Crager, S.E. (2004). What should we be measuring in behavioral studies of chronic pain in animals? Pain 112, 12–15.

27. Langford, D.J., Bailey, A.L., Chanda, M.L., Clarke, S.E., Drummond, T.E., Echols, S., Glick, S., Ingrao, J., Klassen-Ross, T., Lacroix-Fralish, M.L., et al. (2010). Coding of facial expressions of pain in the laboratory mouse. Nat.Methods 7, 447–449.

28. Dolensek, N., Gehrlach, D.A., Klein, A.S., and Gogolla, N. (2020). Facial expressions of emotion states and their neuronal correlates in mice. Science 368, 89–94.

29. Ambrée, O., Richter, H., Sachser, N., Lewejohann, L., Dere, E., de Souza Silva, M.A., Herring, A., Keyvani, K., Paulus, W., and Schäbitz, W.-R. (2009). Levodopa ameliorates learning and memory deficits in a murine model of Alzheimer’s disease. Neurobiology of Aging 30, 1192–1204.

30. Guzmán-Ramos, K., Moreno-Castilla, P., Castro-Cruz, M., McGaugh, J.L., Martínez-Coria, H., LaFerla, F.M., and Bermúdez-Rattoni, F. (2012). Restoration of dopamine release deficits during object recognition memory acquisition attenuates cognitive impairment in a triple transgenic mice model of Alzheimer’s disease. Learning & Memory 19, 453–460.

31. Brooks, S.P., and Dunnett, S.B. (2009). Tests to assess motor phenotype in mice: a user’s guide. Nat.Rev.Neurosci 10, 519–529.

32. Reeves, R.H., Irving, N.G., Moran, T.H., Wohn, A., Kitt, C., Sisodia, S.S., Schmidt, C., Bronson, R.T., and Davisson, M.T. (1995). A mouse model for Down syndrome exhibits learning and behaviour deficits. Nature Genetics 11, 177–184.

33. Mientjes, E.J., Nieuwenhuizen, I., Kirkpatrick, L., Zu, T., Hoogeveen-Westerveld, M., Severijnen, L., Rife, M., Willemsen, R., Nelson, D.L., and Oostra, B.A. (2006). The generation of a conditional Fmr1 knock out mouse model to study Fmrp function in vivo. Neurobiol Dis 21, 549–555.

34. Francardo, V., Recchia, A., Popovic, N., Andersson, D., Nissbrandt, H., and Cenci, M.A. (2011). Impact of the lesion procedure on the profiles of motor impairment and molecular responsiveness to L-DOPA in the 6-hydroxydopamine mouse model of Parkinson’s disease. Neurobiology of Disease 42, 327–340.

35. Stein, C., Millan, M.J., and Herz, A. (1988). Unilateral inflammation of the hindpaw in rats as a model of prolonged noxious stimulation: Alterations in behavior and nociceptive thresholds. Pharmacology Biochemistry and Behavior 31, 445–451.

36. Hazan, L., Zugaro, M., and Buzsaki, G. (2006). Klusters, NeuroScope, NDManager: a free software suite for neurophysiological data processing and visualization. Journal of Neuroscience methods 155, 207–216.

37. Mills, T.C. (1991). Time Series Techniques for Economists, (Cambridge University Press).

38. Hartigan, J.A., and Wong, M.A. (1979). Algorithm AS 136: A k-means clustering algorithm. Applied Statistics 28, 100–108.

39. Cortes, C., and Vapnik, V. (1995). Support-Vector Networks. Machine Learning 20, 273–297.

40. Rijsbergen, C.J.V. (1979). Information Retrieval, (Butterworth-Heinemann).

41. Kriegeskorte, N., Simmons, W.K., Bellgowan, P.S.F., and Baker, C.I. (2009). Circular analysis in systems neuroscience – the dangers of double dipping. Nature neuroscience 12, 535–540.

42. Razali, N.M., and Bee Wah, Y. (2011). Power comparisons of Shapiro-Wilk, Kolmogorov-Smirnov, Lilliefors and Anderson-Darling tests. Journal of Statistical Modeling and Analytics 2, 21–33.

